# What Determines the Success of Restoring the Threatened African Savanna Mahogany (*Khaya senegalensis*) Beyond Seed-Tree Size? A Multi-Model Test of Site and Maternal-Origin Effects

**DOI:** 10.64898/2026.08.12.744298

**Authors:** Beda Innocent Adji, Doffou Sélastique Akaffou

## Abstract

*Khaya senegalensis* (Desr.) A. Juss., the West African savanna mahogany, is listed as Vulnerable on the IUCN Red List due to prolonged commercial and medicinal overexploitation, compounded by failing natural regeneration amid growing deforestation and climate change. Restoration strategies generally rest on the implicit assumption that rigorous selection of seed trees based on their size improves offspring quality. Yet this assumption has never been confronted with the competing hypothesis of a determinism dominated by the planting site, using statistical approaches of adequate power. This study tests the relative contribution of seed-tree dendrometric characteristics (diameter, height) and planting site to germination and juvenile growth (6-24 months) of *K. senegalensis*, using six complementary analytical approaches: continuous regression, linear mixed models with site × seed-tree interaction, seed-mass mediation analysis, non-linear growth-trajectory modelling, between-site phenotypic plasticity indices, and multi-predictor allometric equations, across two contrasting bioclimatic sites in Côte d’Ivoire (Korhogo: Sudanian savanna, and Daloa: humid forest) and six seed-tree categories. Converging across all these approaches, the results show that the planting site exerts a decisive influence on germination (71.1% at Korhogo versus 63.9% at Daloa; χ*²* = 6.09; *p* = 0.014) and on the entire growth trajectory (*p* < 0.001), whereas no seed-tree characteristic affects offspring performance, whether in continuous regression (*p* > 0.37), mixed models with interaction (*p* > 0.29), or through a seed-mass mediation pathway (Sobel test, *p* = 0.46). An exponential growth model fits better than linear or power-law models (Δ*AIC* = 13.6-14.7 depending on site), and multi-predictor allometric equations (diameter and height) explain 97.4% of the variance in total biomass, versus 94.8% for the single-predictor model. These convergent results indicate that, contrary to the widespread practice of selecting seed trees on size criteria, it is the choice of planting site that truly determines the success of restoring this vulnerable species. A conclusion calling for a reorientation of priorities in reforestation and agroforestry programmes across West Africa.

## 1. Introduction

African tropical forests, exceptional reservoirs of plant genetic diversity, are eroding under the combined effect of deforestation, unsustainable agricultural expansion, and climate change (Choat et al., 2012; De Wasseige C et al., 2012; Sosef et al., 2017). In Côte d’Ivoire, this dynamic now extends from the dense forests of the south to the savannas of the north, threatening local woody genetic resources (Adji et al., 2023; Akaffou et al., 2021). A situation to which the country has responded by banning the exploitation of several species north of the 8th parallel and by committing to restore 5.000 hectares by 2030 (Adji et al., 2023). This commitment can only be met if reforestation and agroforestry programmes are grounded in solid knowledge of the regeneration biology of target species. Among the priority species for this effort, *Khaya senegalensis* (Desr.) A. Juss. (Meliaceae), the African mahogany or caïlcédrat, occupies a particularly important place. This species starkly illustrates the tension between economic value and conservation. A large, majestic tree emblematic of the Sudano-Sahelian and Sudanian savannas, it is prized for its high-quality timber, its extensive use in traditional pharmacopoeia against ailments ranging from malaria to dermatoses, and its role in urban forestry, shade provision, and rural livelihoods (Adji et al., 2020; Kerharo and Bouquet, 1950; Tropiques, 1988). This species has been intensively exploited for nearly two centuries (Bouka Dipelet et al., 2019), to the point that it is now listed as Vulnerable (VU) on the IUCN Red List (IUCN, 1998). In Côte d’Ivoire, its natural stands are becoming increasingly scarce, weakened by overexploitation, medicinal harvesting (Doffou et al., 2024; Issa et al., 2018; Kadiri et al., 2025; Silue et al., 2021), attacks by the shoot borer *Hypsipyla robusta*, and a changing climate (Sokpon and Ouinsavi, 2004). This regeneration failure is not unique to *K. senegalensis*, as it echoes a broader pattern documented across the genus *Khaya* and other West African mahoganies, whose taxonomic and ecological boundaries remain only partially resolved despite two centuries of commercial exploitation, complicating the design of species-specific conservation strategies (Bouka Dipelet et al., 2019). In *K. senegalensis* specifically, natural regeneration in the field is characterised by very limited seedling recruitment despite abundant seed production, raising the question of whether the bottleneck lies in the biology of the seed and seedling itself, in the environmental conditions encountered during establishment, or in an interaction between the two. Answering this question through controlled nursery trials (rather than through in situ observation alone) is essential to guide the seed-harvesting and site-selection strategies underpinning large-scale reforestation and agroforestry programmes, and to identify effective levers for restoration.

Two non-mutually exclusive hypotheses have been put forward in the literature to explain the variability in germination success and seedling vigour in *K. senegalensis* and related species. The first attributes this variability to a maternal effect: the dendrometric characteristics of the seed tree (its diameter, height, vigour) could influence the quality, and in particular the mass, of the seeds it produces, with cascading consequences for germination and initial seedling performance, as documented for seed mass across hundreds of tropical tree species (Norden et al., 2009) and, more specifically, for variation in seed size and seedling traits according to provenance and family within *K. senegalensis* itself (Ky-Dembele et al., 2014). The second hypothesis attributes this variability mainly to the germination and growth environment (climate, soil, nursery management), which shapes the expression of the seedling’s genetic potential largely independently of its maternal origin. Adji et al. (2021a, 2021b, 2020) directly tested this second hypothesis by comparing germination and seedling morphology of *Pterocarpus erinaceus*, *Parkia biglobosa*, and *Khaya senegalensis* across three contrasting environments (two Ivorian nurseries and a controlled greenhouse in France), and showed that the environment, but not the dendrometric characteristics of the six sampled seed trees, significantly influenced nearly all germination and growth parameters. This was a striking result, yet one that rested on a categorical seed-tree factor (six discrete ’seed-tree’ groups) rather than on the continuous dendrometric variables themselves, and that did not explore the possibility that a maternal effect might operate indirectly, through seed mass, rather than directly through tree size. This simple categorical group comparison proves very limited, testing neither the continuous relationship with actual tree size nor any potential mediation through seed mass. More recently, building on the same expanded experimental design, Adji et al. (2023) showed that seed size itself (as distinct from seed-tree size) is a strong and consistent predictor of both germination and seedling vigour in *K. senegalensis*. This finding sharpens the central question addressed in this study: if seed size matters so much, does the size of the seed tree that produced it also matter, whether directly or indirectly through its influence on the seed mass it produces?

Disentangling these two levels (the individual seed and the individual seed tree) requires statistical approaches considerably more advanced and structurally finer than the analysis-of-variance framework historically applied to seed-tree comparisons in this species (Adji et al., 2020; Ky-Dembele et al., 2014), namely continuous regression on actual dendrometric measurements, mixed-effects models accounting for the non-independence of seedlings sharing the same mother tree, and a formal mediation analysis capable of testing whether a seed-tree effect, if it exists, is transmitted through seed mass.

The present study addresses this gap by re-examining, in *K. senegalensis*, the relative contribution of seed-tree dendrometric characteristics and site to germination and juvenile (6-24 months) and adult (5 and 7 years) growth, across two contrasting bioclimatic zones of Côte d’Ivoire (the Sudanian savanna of Korhogo and the humid forest zone of Daloa), using six complementary analytical approaches (continuous regression, mixed models, mediation, non-linear growth, phenotypic plasticity, multi-predictor allometry). This study addresses the following questions: (i) does the planting site significantly affect the germination kinetics and juvenile growth trajectories of *K. senegalensis*, and what is the relative magnitude of this effect compared with the site effect already reported for this species (Adji et al., 2020)(ii) do the dendrometric characteristics of the seed tree (diameter and height) exert a direct effect on the performance of its offspring once tested by sufficiently powerful continuous statistical methods, or an indirect effect mediated by seed mass? and (iii) can robust multi-predictor allometric models be established to support non-destructive biomass monitoring in restoration and agroforestry programmes involving this species? We test three hypotheses: (H1) the site exerts a significant and substantial effect on germination and growth; (H2) the seed tree exerts no detectable direct effect once tested by adequate continuous methods; (H3) any residual seed-tree effect is more likely transmitted through seed mass than through its own dimensions. By clarifying this determinism, this study aims to provide forest managers and tree breeders with an unambiguous, statistically robust answer, and to settle a question of direct consequence for the conservation and restoration of this vulnerable species: should seed-harvesting and seed-orchard strategies prioritise the selection of large seed trees, or rather the matching of planting material to favourable planting sites?

## 2. Materials and methods

### 2.1. Study sites

The study was conducted between December 2018 and June 2025, with several repetitions, in two experimental nurseries in Côte d’Ivoire, selected for their marked bioclimatic contrast along a north-south climatic gradient. The Korhogo forestry research station (9°57’0.556" N; 5°54’2.889" W), managed by the Centre National de Recherche Agronomique (CNRA), is representative of the Sudanian zone: dry tropical climate, annual rainfall of 817 to 1216 mm, temperatures of 26.6 to 35.7 °C, shallow, gravelly ferruginous and ferralitic soils, low in organic matter and strongly desaturated (Adji et al., 2023). The Jean Lorougnon Guédé University in Daloa (6°9’6.363" N; 6°43’8.1157" W), in the west-central part of the country, is representative of the sub-equatorial dense humid forest zone: rainfall of 1000 to 1900 mm/year, temperatures of 21 to 34 °C, deep, acidic ferralitic soils, desaturated in exchangeable bases but rich in organic matter (Adji et al., 2023; Akaffou et al., 2021). This pedoclimatic contrast makes it possible to test the site effect independently of the origin of the plant material, which is identical at both sites.

### 2.2. Study species and plant material

*Khaya senegalensis* (Desr.) A. Juss. (Meliaceae) is a tree that can reach 35 m in height, with scaly brown bark, characteristic of Sudano-Guinean savannas and gallery forests (Bouka Dipelet et al., 2019), and listed as Vulnerable on the IUCN Red List (IUCN, 1998). The plant material consists of seeds from 120 seed trees per species, sampled over several harvest campaigns (December 2018, January and February 2019, December 2020, and February 2023) in four localities arranged along a bioclimatic gradient in Côte d’Ivoire: Katiola (5°7’35.814" W; 8°13’53.94" N), Niakara (5°18’40.73544" W; 8°40’47.97912" N), Korhogo (5°36’12.39612" W; 9°33’24.68988" N), and Sinématiali (5°22’59.999" W; 9°34’59.999" N). Seed trees were grouped into six distinct categories according to their dendrometric size. Six reference trees were selected at the *CNRA* forestry research station in Korhogo to represent a gradient of contrasting dendrometric characteristics (diameter at breast height and total height), and 19 additional seed trees with similar characteristics were then associated with each of them from across the four provenance localities mentioned above, forming six groups of 20 seed trees (Table 1). The mean DBH and mean height of each group, calculated over the 20 seed trees, range respectively from 15.1 to 63.2 cm and from 11.6 to 24.7 m (Figure 1); it should be noted that these ranges partially overlap between certain groups, which retrospectively justifies the use, in the analyses presented in section 2.7, of continuous regression and mixed models treating DBH as a real variable rather than a simple categorical comparison, the latter approach being more sensitive to partial overlap between classes. The age of the reference tree in each group, by contrast, was estimated through oral surveys of rural communities and is available only for that single tree, not for all 20 seed trees in the group. Seed trees within the same locality were approximately 200 m apart from one another. Diameter was measured with a forestry tape and height was estimated using the lumberjack’s eye method. The plant material used is the property of rural populations; authorisation to use it for this study was given orally by the orchard owners.

**Fig. 1.**
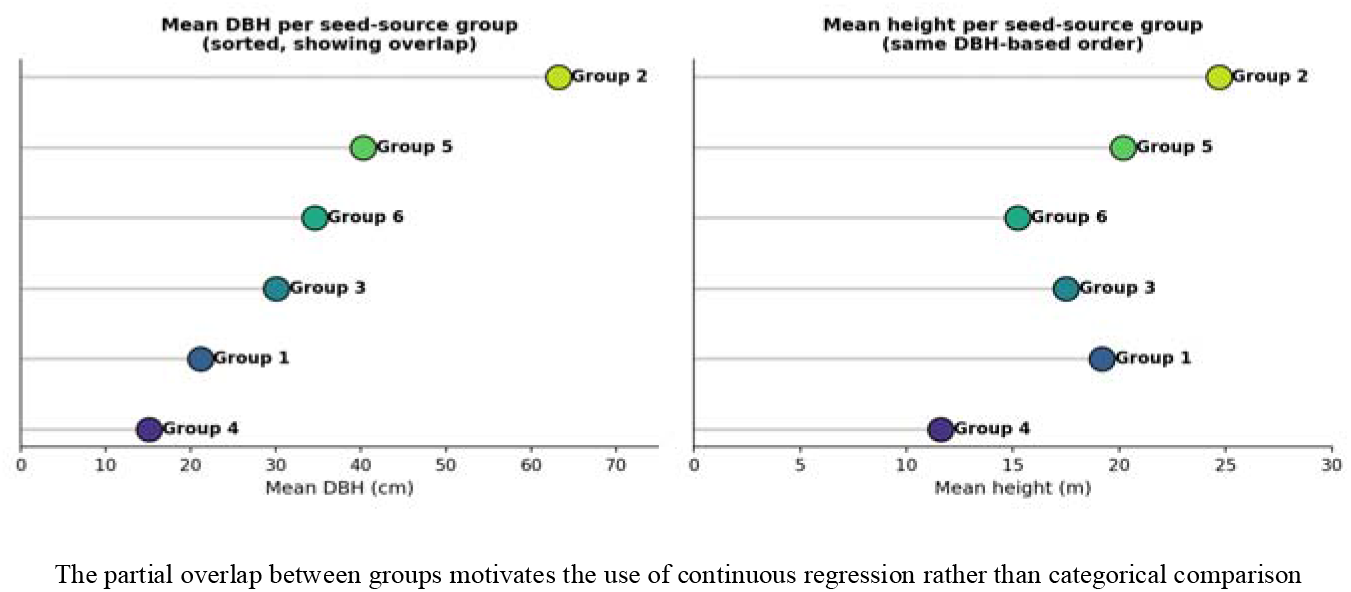
Dendrometric gradient (mean DBH and height) among the six seed-tree groups of *Khaya senegalensis*, sorted by increasing DBH.

**Table 1.** **Dendrometric characteristics of the six seed-tree groups of *Khaya senegalensis***

| Seed-tree group | Mean DBH<sup>a</sup><br>(cm) | Mean H<sup>a</sup><br>(m) | Reference tree age<sup>b</sup><br>(Years) | GPS coordinates of reference tree |  | Number of seed trees | Seed mass (g) |  |  |
| --- | --- | --- | --- | --- | --- | --- | --- | --- | --- |
|  |  |  |  | Longitude | Latitude |  | Min | Max | Mean |
| 1 | 21.15 | 19.2 | 32 | -5.5033 W | 9.5649 N | 20 | 0.1 | 0.27 | 0.16 ± 0.04 |
|  |  |  |  |  |  |  |  |  | ab |
| 2 | 63.2 | 24.7 | 43 | -5.54454 W | 9.56562 N | 20 | 0.06 | 0.26 | 0.18 ± 0.05 |
|  |  |  |  |  |  |  |  |  | ab |
| 3 | 30.06 | 17.5 | 27 | -5.54824 W | 9.56603 N | 20 | 0.06 | 0.32 | 0.21 ± 0.07 |
|  |  |  |  |  |  |  |  |  | a |
| 4 | 15.11 | 11.61 | 28 | -5.55008 W | 9.56476 N | 20 | 0.03 | 0.19 | 0.10 ± 0.04 |
|  |  |  |  |  |  |  |  |  | b |
| 5 | 40.29 | 20.18 | 32 | -5.5051 W | 9.56498 N | 20 | 0.09 | 0.28 | 0.17 ± 0.05 |
|  |  |  |  |  |  |  |  |  | ab |
| 6 | 34.55 | 15.23 | 32 | -5.55064 W | 9.56345 N | 20 | 0.15 | 0.46 | 0.26 ± 0.08 |
|  |  |  |  |  |  |  |  |  | a |
| **Pr > F** |  |  |  |  |  |  |  |  | **0.014** |

Mean DBH = Mean diameter at breast height of the tree groups in centimetres; Mean H = Mean height of the tree groups in metres; W = West; N = North; Min = Minimum; Max = Maximum. <sup>a</sup>Mean calculated across the 20 seed trees of the group (diameter measured with a forestry tape; height estimated using the lumberjack's eye method). <sup>b</sup>Age estimated through oral surveys of rural communities, reported only for the reference tree used to define each group (not available for all 20 seed trees). Since DBH and height served as the criteria for constituting the groups, no inter-group statistical test was applied to them; only seed mass (the outcome variable) was tested by ANOVA (values sharing the same letter: no significant difference, $p < 0.05$ ).

Seedlings originating from the germination of seeds from the six seed-tree categories were monitored at different ages (4 months, 6 months, 1 year, 2 years, 5 years, and 7 years). Seeds destined for germination and growth trials were extracted manually, sorted by flotation, and air-dried until moisture stabilised (4.5-5%), following the ISTA (2009) protocol. The present study specifically uses the germination and growth data collected at 6, 12, and 24 months at the two contrasting sites of Korhogo and Daloa, constituting a subset of the broader experimental design described above; monitoring at 4 months, 5 years, and 7 years, as well as the two other seed-collection localities (Katiola, Niakara), are not covered by the analyses presented here (no provenance test).

### 2.3. Experimental design

Seeds from each seed-tree group were divided into two lots destined respectively for Korhogo and Daloa. After 12 hours of soaking to break seed-coat dormancy, seeds were sown at a depth of 1.5-2 cm (with the lower end of the seed directed into the soil and the upper, wider end towards the surface) in polyethylene bags (20 × 10 cm) filled with local potting soil, at a rate of two seeds per bag, arranged in a block subdivided into six sub-blocks representing each seed-tree group. Each sub-block, containing 60 bags, was labelled with the serial number of the seed-tree group (giving 2 seeds × 60 pots × 6 groups × 2 sites = 1.440 seeds for the trial). Seeds were collected and sown in subsequent years as experimental repetition and replacement. A phytosanitary treatment (granular FURADAN against rodents, DECIS against larval attacks) was applied, and maintenance consisted of daily watering and manual weeding.

### 2.4. Germination monitoring

Germination, defined by the appearance of the hypocotyl at the substrate surface, was recorded daily at each site, allowing a cumulative germination model to be fitted and five classical parameters to be calculated (lag time, delay, rate, duration, and germination percentage), following the definitions of Adji et al. (2023, 2020).

### 2.5. Growth and biomass monitoring

After germination, 30 vigorous seedlings were randomly selected per seed-tree group and per site for non-destructive growth monitoring at 6, 12, and 24 months: total height (from collar to apex), collar diameter, number and dimensions of leaves, root length and diameter, measured using a graduated ruler, electronic calipers, and ImageJ software for leaf area. A subsample of plants was harvested at the same ages for destructive biomass measurement (total, foliar, stem, and root fresh and dry mass), after oven-drying (70 °C, 72 h) and weighing to the nearest 0.001 g, allowing allometric equations to be established.

### 2.6. Data quality control

A systematic consistency check was applied to the entire dataset prior to analysis, verifying in particular that the dry root mass of each individual did not exceed its total dry mass. Detected outliers (5.9%) were treated as missing data rather than arbitrarily corrected, and explicitly documented. This step, rarely explicitly reported in the literature on the regeneration of African forest tree species, is presented here in the interest of reproducibility and scientific transparency.

### 2.7. Analytical strategy

In contrast to the approach of comparing seed-tree categories by classical analysis of variance, historically used for *K. senegalensis* (Adji et al., 2020) and other related species (Ky-Dembele et al., 2014), six complementary statistical approaches were used to maximise the power to detect any seed-tree effect and to clearly distinguish it from the site effect:

#### 2.7.1. Continuous regression

Rather than comparing seed-tree groups against one another, as if tree size were a fixed category, we directly tested whether the actual diameter (DBH) and height of the seed tree can predict the performance of its offspring (much as one would test whether parents’ height predicts that of their children, by plotting a scatter plot and fitting a line through it). Specifically, each growth parameter was regressed on DBH and then on seed-tree height, separately at each age, by simple linear regression. For each growth parameter *Y* (height, collar diameter, total biomass) and at each measurement age, a simple linear regression model was fitted separately on the actual DBH and height of the original seed tree, rather than on a categorical group variable:

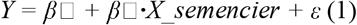

*Y* is the growth parameter, *X*_seed-tree is the seed tree’s DBH or height, and β□ is the tested coefficient (fitted by ordinary least squares), or more explicitly, *Y* is the growth parameter considered, *X*_seed-tree alternately represents the seed tree’s DBH or height (continuous variable, in cm or m), β□ is the intercept, β□ is the tested regression coefficient (*H0*: β□ = 0), and ε is the residual error term, assumed i.i.d. *∼ N*(0, σ*²*). The model was fitted by ordinary least squares (*OLS*), separately for each parameter × age × predictor combination (18 regressions in total: 3 parameters × 3 ages × 2 predictors).

#### 2.7.2. Linear mixed models

For each response *Y* (height, diameter, log-biomass), a linear mixed model was fitted with seed tree as a random effect (accounting for the non-independence of individuals originating from the same mother tree) and site, age, and seed-tree DBH as fixed effects, including a DBH × site interaction term. Indeed, seedlings from the same seed tree are not entirely independent of one another: they share a common origin, much like siblings within the same family. Ignoring this relatedness can bias classical statistical tests. The mixed model corrects for this by treating each seed tree as a ’family’ whose own variability is estimated, while simultaneously testing the effect of DBH, site, and their interaction (is the effect of DBH different at Daloa and Korhogo?) on growth:

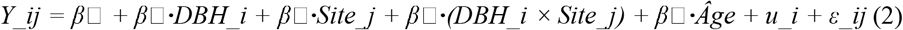

where *Y_ij* is the observed value for the individual from seed tree *i* at site *j*; *u_i* ∼ *N*(0, σ*²_u*) is the random effect of seed tree i (accounting for between-seed-tree variance); ε*_ij ∼ N*(0, σ*²_*ε) is the individual-level residual. Coefficient β□ tests the main (direct) effect of seed-tree DBH; β□ tests whether this effect differs by site (interaction). In other words, *u_i* is the effect specific to seed tree *i* (random effect) and ε*_ij* is the individual residual. β□ tests the direct effect of DBH; β□ tests its interaction with site. Models were fitted by restricted maximum likelihood (REML; Bates et al., 2015), using the lme4/lmerTest packages (*R*). For biomass, the response variable was log-transformed (’log(total dry mass)’) to stabilise variance, as the raw distribution was strongly right-skewed.

#### 2.7.3. Mediation analysis and Sobel test

Even if seed-tree DBH has no direct effect on the growth of its offspring, it could have a hidden, indirect effect through an intermediate step: a large tree might produce heavier seeds, and these heavier seeds might in turn germinate better and produce more vigorous seedlings. We tested this pathway (DBH → seed mass → germination → growth) in several steps at the level of the six seed-tree groups (*n* = 6) through three simple, successive sequential regressions (paths *a* to *c*), each testing one link in the chain (Baron and Kenny, 1986) :

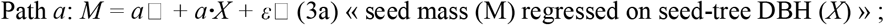

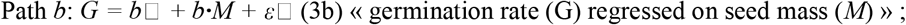

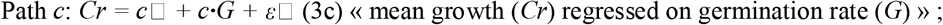

Total effect: *Cr = c’*□ *+ c’**·**X +* ε□ « growth regressed directly on seed-tree DBH, with no mediator » and the indirect effect of *X* on *G* via *M* is estimated by the product of the coefficients: Indirect effect = *a × b*. A complementary test (the Sobel test) allows verification of whether the overall indirect effect (through all links) is statistically significant, even if each link taken in isolation is not fully significant:

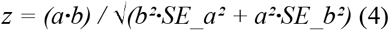

where *SE_a* and *SE_b* are the standard errors of coefficients *a* and *b* respectively. The *z* statistic is compared against the standard normal distribution (two-tailed test, α = 0.05 threshold).

#### 2.7.4. Non-linear modelling of growth trajectories

Plants do not necessarily grow at a constant rate over time: some species grow slowly at first then accelerate, while others slow down as they age. We therefore compared three possible shapes of height-growth curve (a straight line, « steady growth »; a power-law curve, « progressively slowing growth »; and an exponential curve, « growth that accelerates over time ») to identify which best describes the actual seedling trajectory, separately for each site. The choice among these three curves was made using Akaike’s Information Criterion (*AIC*), an index that rewards the model offering the best trade-off between goodness of fit and simplicity (an unnecessarily complex model is penalised even if it fits the data slightly better). Three competing models were fitted to the individual height data (*H*) as a function of age (t, in months), separately by site, by non-linear least-squares regression (Levenberg-Marquardt) :

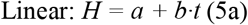

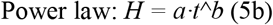

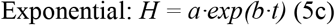

The three models were compared using Akaike’s Information Criterion, calculated from the residual sum of squares (*RSS*) on a common scale (raw, untransformed scale) to ensure a fair comparison between models:

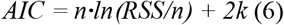

where n is the number of observations and k is the number of model parameters (*k* = 2 for the three models tested). The retained model is the one with the lowest *AIC* (Δ*AIC* > 2 being considered a substantial difference in support between models (Burnham and Anderson, 2004, 2002; Paine et al., 2012).

#### 2.7.5. Phenotypic plasticity index

Beyond mean performance, we sought to determine whether certain seed trees produce offspring that are more ’sensitive’ to site conditions than others. That is, offspring whose growth varies markedly depending on whether they grow at Daloa or Korhogo (high plasticity), as opposed to offspring that behave similarly at both sites (low plasticity). A between-site phenotypic plasticity index of the RDPI type (*Relative Distance Plasticity Index*, Valladares et al., 2006) was calculated for height, by seed-tree group *i* and age *t*:

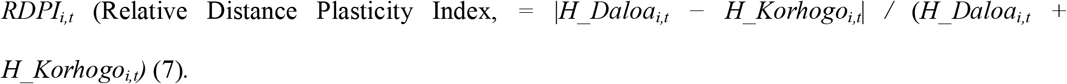

where *H_Daloa_i,t_* and *H_Korhogo_i,t_*are the mean heights of seed-tree group *i* at age *t*, at Daloa and Korhogo respectively. The index ranges from 0 (no plasticity, identical performance at both sites) to 1 (maximum plasticity, if performance differs greatly). The relationship between this index and seed-tree DBH was then tested by regression and by one-way analysis of variance (seed-tree group).

#### 2.7.6. Allometric equations

Measuring the biomass (dry weight) of a plant requires cutting and oven-drying it. A destructive measurement, impossible to carry out across an entire nursery or reforestation trial. Allometric equations circumvent this problem: they relate biomass to simple, non-destructive measurements, such as collar diameter or height, taken on a sample of sacrificed plants, then applied to all other plants without having to cut them down. We compared a model using diameter alone with a model combining diameter and height, to determine which better predicts actual biomass. Total dry biomass (*B*) was modelled as a function of collar diameter (*D*) alone, then of diameter and height (*H*) combined, by log-log regression (power law; Chave et al., 2014; Niklas, 1994) :

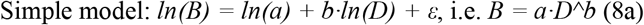

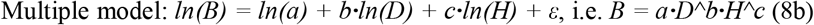

The two models were compared by *AIC* (Equation 6, applied here directly to the regression residuals on the logarithmic scale, both models sharing the same transformed response variable, which makes the comparison directly valid). The retained model is the one that minimises *AIC* while maximising adjusted *R²* (Chave et al., 2014; Niklas, 1994).

All analyses were conducted in *R* and *Python* 3.12 (pandas, numpy, scipy.optimize, statsmodels libraries), with a significance threshold set at α = 0.05 for all tests. The complete scripts (*Python* and *R*) are available and will be provided upon reasonable request to the corresponding author to ensure full reproducibility of the study.

## 3. Results

### 3.1. Germination: overall phenology

Germination of *Khaya senegalensis* is hypogeal, with an epicotyl reaching an average of 6.33 cm in length and 1.2 mm in mean collar diameter (approximately 15 days after sowing and five (5) days after the appearance of the coleoptile and radicle). The prophylls are nearly sessile (petiole absent or stunted), opposite, and generally identical in length and width. The coleoptile and radicle appear on average 10 days after sowing. It is following the prophylls that phyllotaxis becomes alternate-spiral. The seedling gradually produces simple leaves with increasingly long petioles until, at 11 weeks, imparipinnate compound leaves appear, which over time transform into paripinnate compound leaves.

### 3.2. Germination: a clearly marked site effect

The cumulative germination kinetics differ markedly between sites (Figure 2). The final rate reached 71.1 ± 2.0% at Korhogo (standard error, *n*=540 seeds) versus 63.9 ± 2.1% at Daloa (*n*=540 seeds) (χ*²*=6.09; *p*=0.014), with contrasting dynamics: faster emergence at Daloa (*T50*=18.4 days) but a lower final rate than at Korhogo (*T50*=22.6 days). Germination proceeds faster at Daloa early in the cycle (*T50* = 18.4 days) than at Korhogo (*T50* = 22.6 days), but plateaus at a lower level. This result suggests that while Daloa’s conditions favour early emergence, those at Korhogo ultimately allow a larger proportion of the seed lot to germinate.

**Fig. 2.**
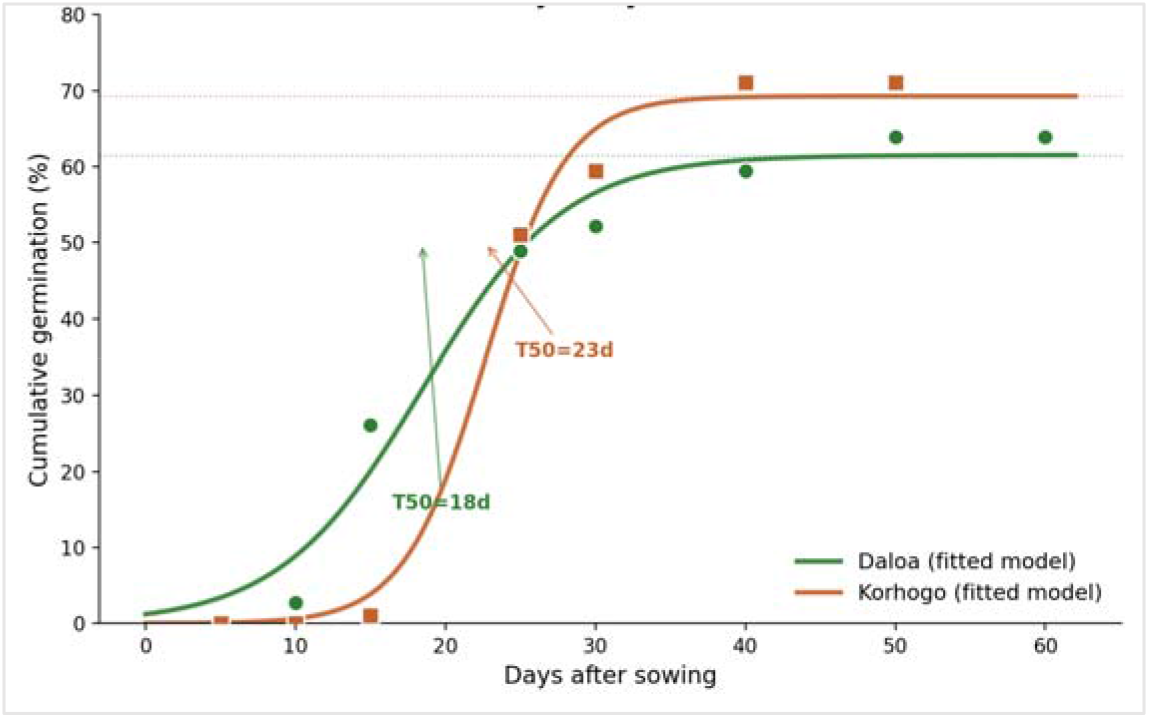
Cumulative germination kinetics of *Khaya senegalensis* by study site, with fitted logistic models (*Gmax* = asymptotic germination rate; *T50* = time to reach 50% of final germination).

Analysis of variance (Table 2) indicates a significant difference between study sites for lag time, germination delay, and germination spread (*P*<0.05). Germination rate and germination speed parameters were statistically similar (*P*>0.05). However, this analysis of variance for each of the six seed-tree groups (all sites combined) shows that seed-tree type has no significant influence on seed germination parameters (*P*>0.05). These germination parameters (Table 2), from the main six seed-tree-group design, are independent of the cumulative kinetics presented in Figure 2, which relies on a distinct sub-sample used to characterise daily germination dynamics across all seed sizes combined.

**Table 2.** Overall influence of study sites and seed-tree groups on germination.

| Sites/Seed-tree group | Lag time | Germination delay | Germination rate | Germination duration | Germination percentage |
| --- | --- | --- | --- | --- | --- |
| **Korhogo** | 21.33 ± 0.21 **a** | 31.74 ± 1.03 **a** | 37.50 ± 2.26 **a** | 23.33 ± 1.94 **a** | 75.22 ± 7.22 **a** |
| **Daloa** | 14.33 ± 1.16 **b** | 23.08 ± 2.07 **b** | 34.16 ± 4.26 **a** | 19.83 ± 3.70 **b** | 76.72 ± 5.20 **a** |
| ***Pr > F*** | **0.0031** | **0.001** | **0.055** | **0.016** | **0.3343** |
| **Group-1** | 14.33 ± 3.38 **a** | 20.49 ± 3.80 **a** | 28.33 ± 3.48 **a** | 16.66 ± 2.90 **a** | 76.77 ± 13.51 **a** |
| **Group-2** | 17.33 ± 2.90 **a** | 25.87 ± 4.78 **a** | 36.66 ± 9.83 **a** | 20.66 ± 6.01 **a** | 72.11 ± 8.81 **a** |
| **Group-3** | 16.00 ± 3.00 **a** | 25.50 ± 5.15 **a** | 30.00 ± 8.00 **a** | 19.66 ± 6.36 **a** | 83.33 ± 8.82 **a** |
| **Group-4** | 17.33 ± 1.85 **a** | 26.09 ± 4.03 **a** | 34.00 ± 7.64 **a** | 21.66 ± 6.36 **a** | 77.21 ± 10.01 **a** |
| **Group-5** | 15.00 ± 3.21 **a** | 21.88 ± 3.96 **a** | 28.00 ± 6.43 **a** | 15.00 ± 2.64 **a** | 73.88 ± 12.37 **a** |
| **Group-6** | 16.66 ± 2.33 **a** | 23.74 ± 5.27 **a** | 25.66 ± 4.97 **a** | 14.33 ± 6.33 **a** | 98.33 ± 1.66 **a** |
| ***Pr > F*** | **0.9598** | **0.9255** | **0.8775** | **0.8848** | **0.4908** |

For each trait, values sharing the same letters are not statistically different at the 5% threshold.

### 3.3. Growth: site outweighs seed-tree type

#### 3.3.1. Morphology at four months of age

The results in Table 3 present the developmental parameters at 4 months by site and seed lot. These results show that seedling morphological parameters are statistically different from one site to another (*P*<0.05), with the exception of leaf length and width, which are statistically identical regardless of study site (*P*<0.05). The overall analysis of variance (all sites combined) of seedling morphological characteristics according to seed-tree group (Table 3) indicates a statistical similarity among the variables Height (*HtPl*), collar diameter (*Dcol*), number of leaves (*NbreFe*), and internode length (*LgEntr*) of seedlings regardless of seed-tree type (*P*>0.05). By contrast, leaf length (*LgFe*) and width (*LaFe*) both differ significantly from one seed-tree group to another (*P<0.05*).

**Table 3.** Overall influence of study sites and seed-tree groups on seedling development.

| Sites/Seed-tree group | *HtPl* (cm) | *Dcol* (mm) | *NbreFe* | *LgFe* (cm) | *LaFe* (cm) | *LgEntr* (cm) |
| --- | --- | --- | --- | --- | --- | --- |
| **Korhogo** | 24.95±0.90 **a** | 3.35±0.22 **a** | 8.33±0.20 **a** | 11.19±0.73 **a** | 3.68±0.18 **a** | 3.22±0.24 **a** |
| **Daloa** | 18.02±0.39 **b** | 2.35±0.22 **b** | 6.33±0.20 **b** | 10.94±0.67 **a** | 3.54±0.19 **a** | 2.81±0.38 **b** |
| ***Pr > F*** | **0.0301** | **0.022** | **0.017** | **0.8310** | **0.4829** | **0.0481** |
| **Group-1** | 22.83±3.04 **a** | 3.01±0.57 **a** | 7.52±0.88 **a** | 10.08±0.26 **b** | 3.17±0.21 **b** | 3.05±0.53 **a** |
| **Group-2** | 22.41±2.98 **a** | 3.73±0.57 **a** | 7.51±0.88 **a** | 11.12±0.18 **b** | 3.96±0.17 **ab** | 3.42±0.46 **a** |
| **Group-3** | 24.68±3.05 **a** | 3.65±0.58 **a** | 8.23±0.88 **a** | 13.29±0.56 **a** | 4.23±0.22 **a** | 3.64±0.21 **a** |
| **Group-4** | 23.62±3.35 **a** | 2.41±0.57 **a** | 7.61±0.88 **a** | 9.94±0.66 **b** | 3.53±0.11 **ab** | 3.22±0.18 **a** |
| **Group-5** | 23.48±2.94 **a** | 3.85±0.57 **a** | 8.47±0.88 **a** | 9.60±0.55 **b** | 3.69±0.17 **ab** | 2.80±0.67 **a** |
| **Group-6** | 24.99±2.93 **a** | 3.42±0.57 **a** | 8.62±0.88 **a** | 11.54±0.47 **b** | 3.04±0.31 **ab** | 3.66±0.68 **a** |
| ***Pr > F*** | **0.9882** | **0.5071** | **0.8892** | **0.0014** | **0.0441** | **0.7933** |

For each trait, values sharing the same letters are not statistically different at the 5% threshold. *HtPl*: Seedling height; *Dcol*: Seedling collar diameter; *NbreFe*:
Number of leaves; *LgFe*: Leaf length; *LaFe*: Leaf width; *LgEntr*: Internode length; *cm*: centimetres; *mm*: millimetres.

#### 3.3.2. Growth and modelling at 6, 12, and 24 months

##### Site, not seed-tree type, determines growth

Three independent and convergent statistical approaches demonstrate that the growth of *K. senegalensis* is determined by the planting site, and not by the dendrometric characteristics of the original seed tree.

##### a) Analysis of variance and box plots

A classical ANOVA confirms and quantifies the magnitude of the site effect on three growth parameters (height, collar diameter, and total dry biomass) at each age (Table 4; Figure 3). Means are systematically reported with their standard deviation.

**Fig. 3.**
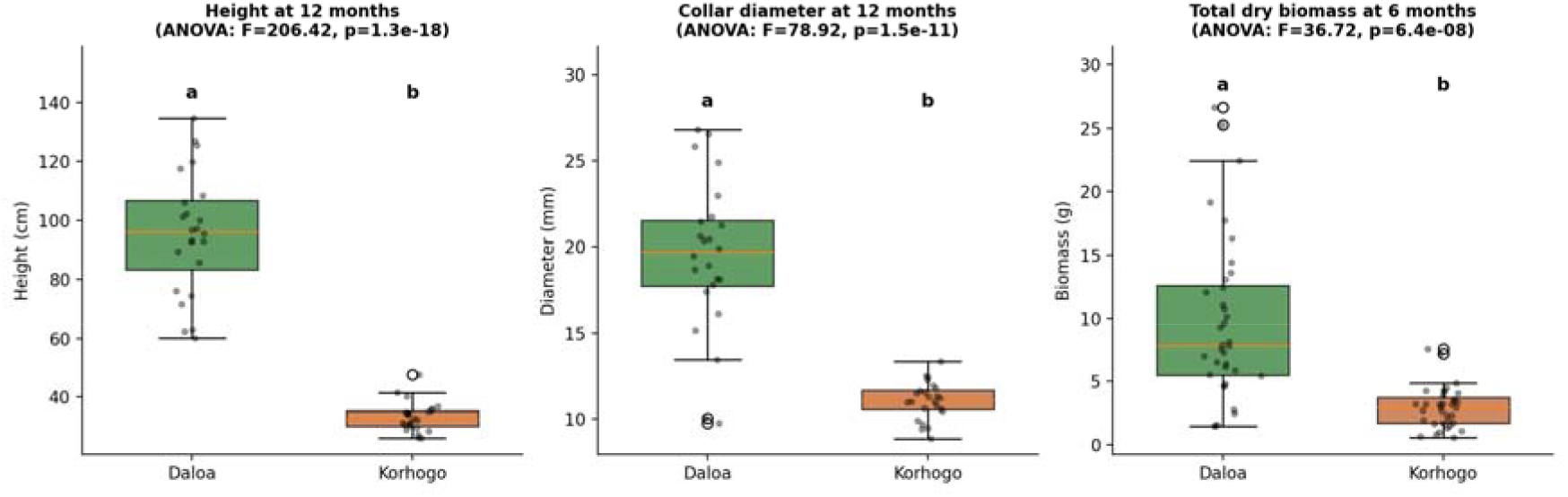
Boxplots showing the effect of study site on height, collar diameter, and total dry biomass of *Khaya senegalensis*, with ANOVA results (letters denote significant differences, *p*<0.05)

**Table 4.** Mean ± standard deviation of the main growth parameters of Khaya senegalensis by site and age.

| Age | Parameter | Daloa<br>(mean±SD) | Korhogo<br>(mean±SD) | *p* |
| --- | --- | --- | --- | --- |
| 6 months | Height (cm) | 49.6±14.9 | 22.9±4.7 | <0.001 |
| 6 months | Biomass (g) | 9.71±6.42 | 2.94±1.62 | <0.001 |
| 12 months | Height (cm) | 95.6±20.7 | 33.3±5.0 | <0.001 |
| 12 months | Biomass (g) | 107.3±53.5 | 15.9±3.5 | <0.001 |
| 24 months | Height (cm) | 424.6±81.0 | 180.1±43.5 | <0.001 |
| 24 months | Biomass (g) | 4757.9±4798.0 | 126.6±57.2 | <0.001 |

For comparison, Figure 4 presents the same type of analysis for the effect of seed-tree group on height at 12 months: no significant difference appears among the six groups, in contrast to the massive site effect observed above.

**Fig. 4.**
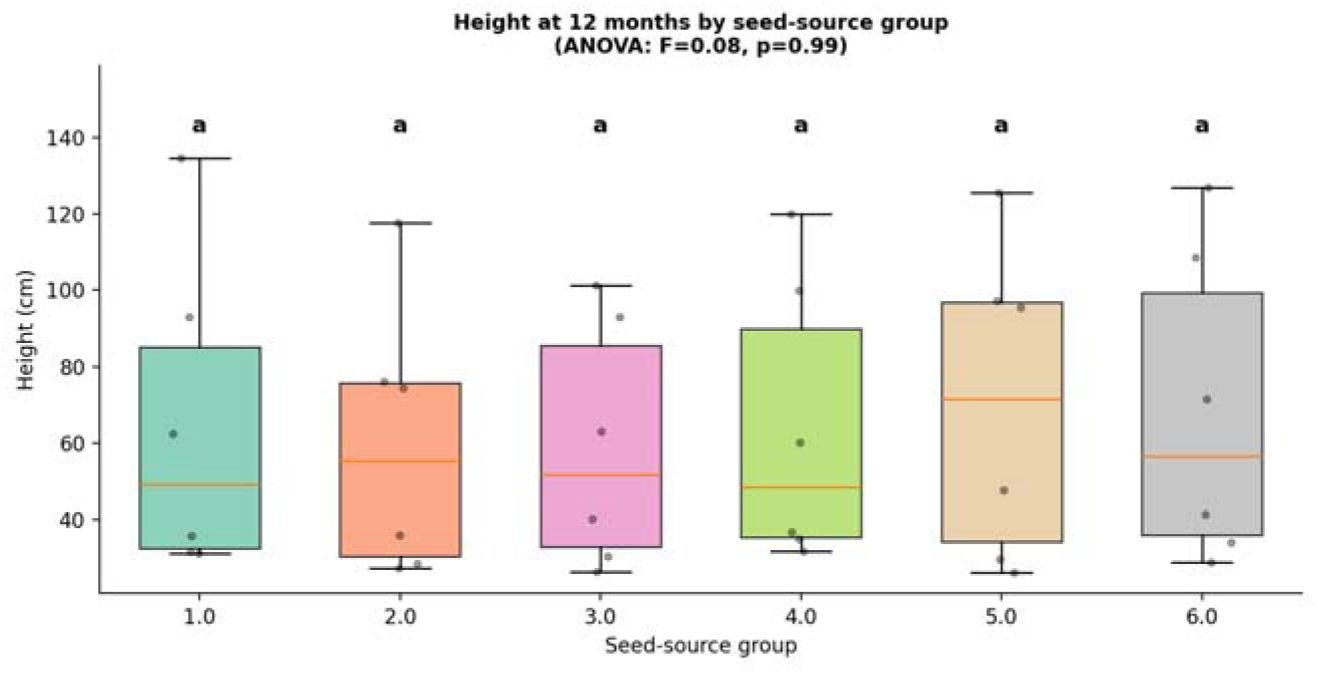
Height at 12 months by seed-source group: no significant difference between mother-tree groups (boxplot with ANOVA).

##### b) Continuous regression

Rather than comparing seed-tree categories, each growth parameter was regressed directly on the actual DBH and height of the original seed tree, separately at each age (Figure 5). Of the 18 regressions tested (3 parameters × 3 ages × 2 predictors), none reached significance (*p* ranging from 0.37 to 0.91; *R²* < 0.03 in all cases). This total absence of a trend, including for biomass at 24 months where the sample size remains adequate (*n*=22), constitutes a substantially more rigorous statistical proof than the traditional simple categorical comparison.

**Fig. 5.**
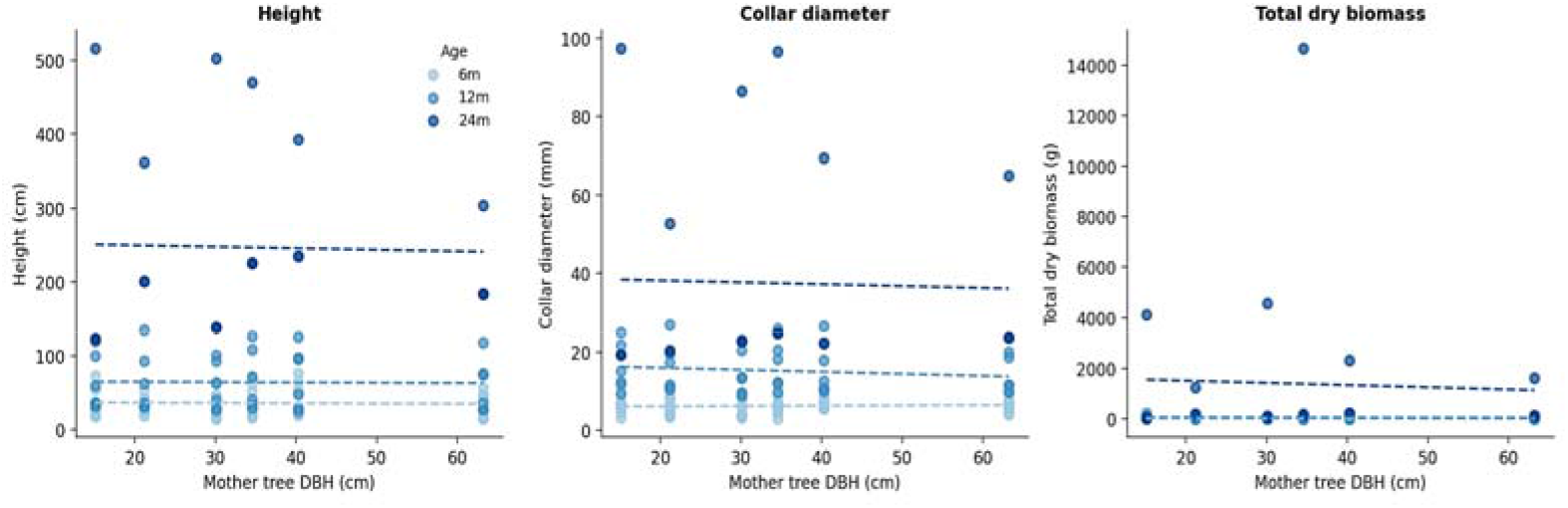
Continuous regression of growth parameters against mother tree DBH, by age class. No relationship is significant (all *p*>0.37).

##### c) Linear mixed models with interaction: site × seed-tree group

Linear mixed models were fitted for height, collar diameter, and total biomass (log-transformed), with seed tree as a random effect, site and age as fixed effects, and a DBH × site interaction term to test whether the seed-tree effect differs according to the environment (Table 5).

**Table 5.** Results of linear mixed models (seed tree as random effect) for three growth responses.

| Response | Seed-tree DBH (*p*) | DBH×site interaction (*p*) | Site (*p*) | Age (*p*) |
| --- | --- | --- | --- | --- |
| **Height** | 0.373 | 0.297 | <0.001 | <0.001 |
| **Collar diameter** | 0.465 | 0.461 | 0.002 | <0.001 |
| **Total biomass (log)** | 0.664 | 0.378 | <0.001 | <0.001 |

Neither the main effect of seed-tree DBH nor its interaction with site is significant for any of the three responses. In other words, the absence of a seed-tree effect is not masked by a differential effect depending on site: it is robust in both environments.

##### c) Classical analysis of variance (site × seed-tree group × age)

In addition to the three preceding approaches, a classical analysis of variance (Table 6, Figures 6 and 7) with main effects (site, seed-tree group, age, without a saturated interaction) was conducted for three growth responses, reproducing the analytical format usually used in this type of study. This classical ANOVA fully confirms the results of the previous approaches: site and age are highly significant for all three responses (*p*<0.001, Table 6, Figures 6 and 7), whereas seed-tree group never is (*p*=0.49 to 0.96, Table 6, Figures 6 and 7). A saturated model including all triple interactions was also tested but proved over-parameterised given the sample size (some *site×group×age* cells contained only a single individual), producing uninterpretable significance artefacts. The more parsimonious main-effects model was therefore retained as the reference, consistent with the convergent results of the continuous regressions and mixed models. The classical ANOVA used here is merely a complementary robustness check rather than a 4th standalone approach.

**Fig. 6.**
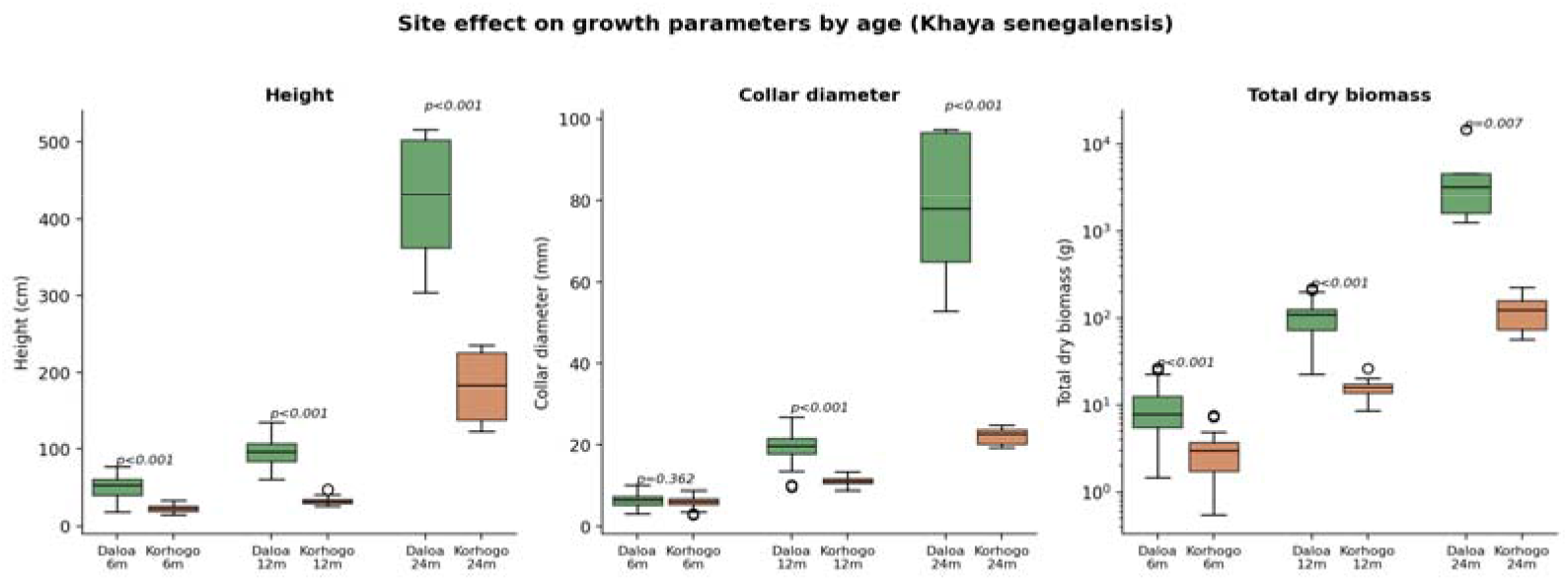
Boxplots of growth parameters (height, collar diameter, total dry biomass) by site and age class, with *Welch’s t-test p*-values.

**Fig. 7.**
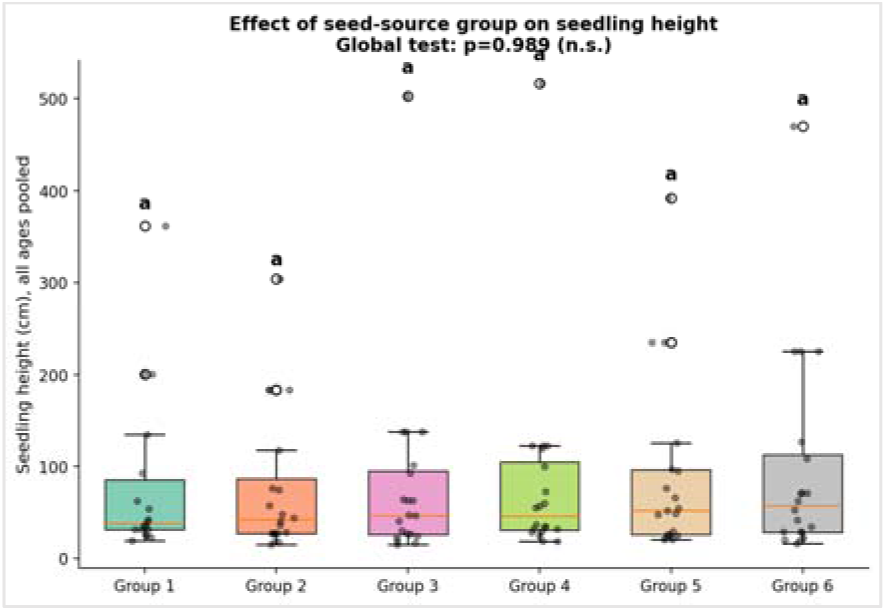
Boxplot of seedling height by seed-source group (all ages pooled), with individual data points. Global *ANOVA* test: *p*=0.989 (n.s.); identical letters indicate no significant difference among groups.

**Table 6.** Analysis of variance with main effects (site, seed-tree group, age) for three growth responses of *Khaya senegalensis*.

| Response | Site (*F*; *p*) | Seed-tree group (*F*; *p*) | Age (*F*; *p*) |
| --- | --- | --- | --- |
| Height | 65.78 ; <0.001 | 0.68 ; 0.638 | 189.25 ; <0.001 |
| Collar diameter | 31.67 ; <0.001 | 0.20 ; 0.963 | 75.77 ; <0.001 |
| Total biomass (log) | 136.73 ; <0.001 | 0.90 ; 0.487 | 310.42 ; <0.001 |

##### d) Mediation analysis (no indirect effect via seed mass): does the seed tree act through seed mass?

An alternative hypothesis is that seed-tree size could indirectly influence the performance of its offspring through the mass of the seeds it produces (a classical maternal effect in seed ecology). A mediation pathway was tested at the level of the six seed-tree groups: seed-tree DBH → mean seed mass → germination rate → mean seedling height (Figure 8).

**Fig. 8.**
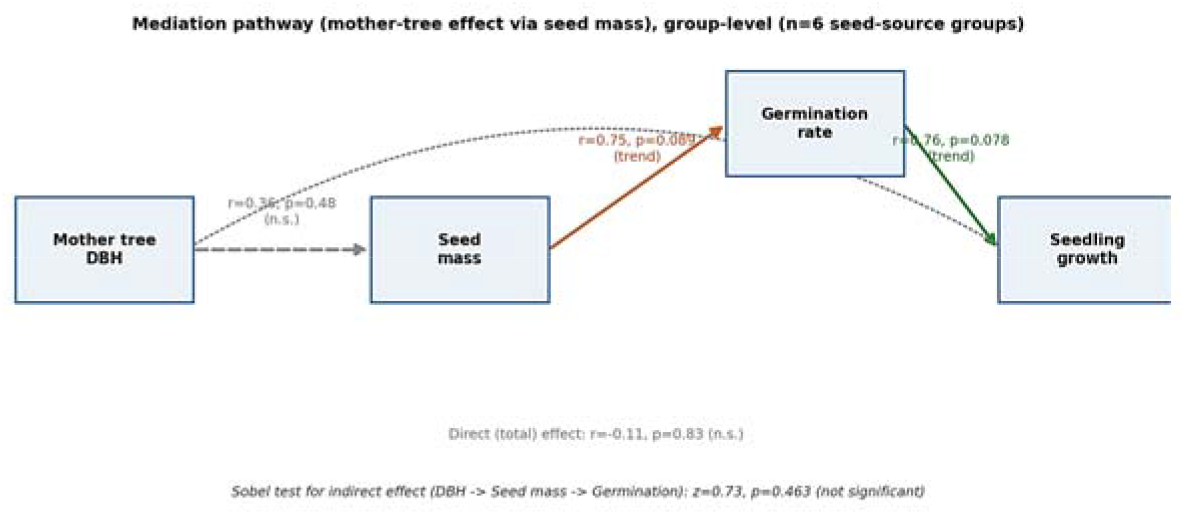
Mediation path linking mother tree DBH to offspring growth through seed mass and germination rate (group-level analysis, *n*=6 seed-source groups).

The first link in the causal chain (DBH → seed mass) is not significant (*r*=0.36; *p*=0.48), which breaks the assumed mediation mechanism. The next two links (seed mass → germination, *r*=0.75, *p*=0.089; germination → growth, *r*=0.76, *p*=0.078) show positive trends close to the significance threshold, but with only six seed-tree groups. Statistical power remains low, and these results should be interpreted as exploratory. The Sobel test for the overall indirect effect (DBH → seed mass → germination) is not significant (*z*=0.73; *p*=0.46). This analysis therefore confirms, through a different route, that seed-tree size has no detectable effect on offspring performance, including via the indirect route of seed mass.

##### e) Non-linear modelling of growth trajectories

Three growth models (linear, power law, exponential) were fitted to individual height data as a function of age, separately by site, and compared by *AIC* calculated on a common scale (Tables 7 and 8). A Gompertz model was also tested, but its asymptote parameter proved poorly constrained by the three available age classes (6-24 months), the time window being too short relative to the species’ lifespan; this model was therefore not retained.

**Table 7.** Comparison of height growth models by *AIC* (common scale, see. **Figure 9).**

| Site | Model | AIC | R<sup>2</sup> |
| --- | --- | --- | --- |
| Daloa | Linear | 570.6 | 0.865 |
| Daloa | Power law | 532.2 | 0.921 |
| Daloa | Exponential | 517.5 | 0.935 |
| Korhogo | Linear | 503.4 | 0.821 |
| Korhogo | Power law | 469.9 | 0.885 |
| Korhogo | Exponential | 456.3 | 0.904 |

**Table 8.**
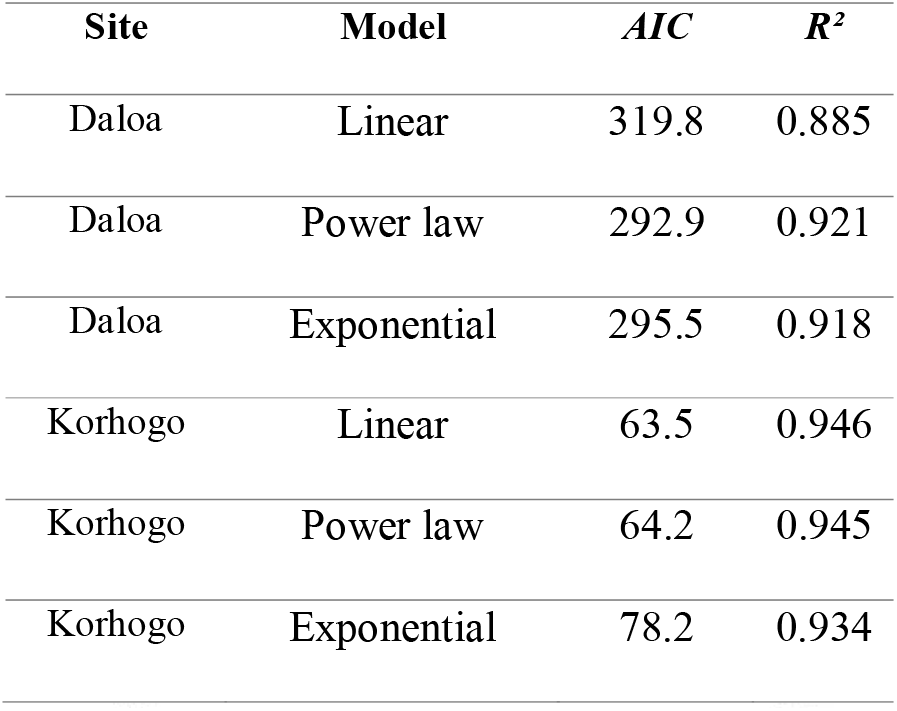
Comparison of collar-diameter growth models by *AIC* (common scale; see. **Figure 10).**

| Site | Model | *AIC* | *R*<sup>2</sup> |
| --- | --- | --- | --- |
| Daloa | Linear | 319.8 | 0.885 |
| Daloa | Power law | 292.9 | 0.921 |
| Daloa | Exponential | 295.5 | 0.918 |
| Korhogo | Linear | 63.5 | 0.946 |
| Korhogo | Power law | 64.2 | 0.945 |
| Korhogo | Exponential | 78.2 | 0.934 |

#### Height growth

The exponential model is consistently the best fit at both sites (Δ*AIC* = 13.6 at Korhogo and 14.7 at Daloa relative to the power law), indicating that no growth slowdown is yet perceptible between 6 and 24 months. The estimated parameters (Daloa: *H*=22.7×*exp*(0.122×*age*); Korhogo: *H*=8.2×*exp*(0.128×*age*)) show very similar relative growth rates between sites, meaning that most of the growth gap stems from a difference in initial height rather than a difference in relative growth rate (Table 7, Figure 9).

**Fig. 9.**
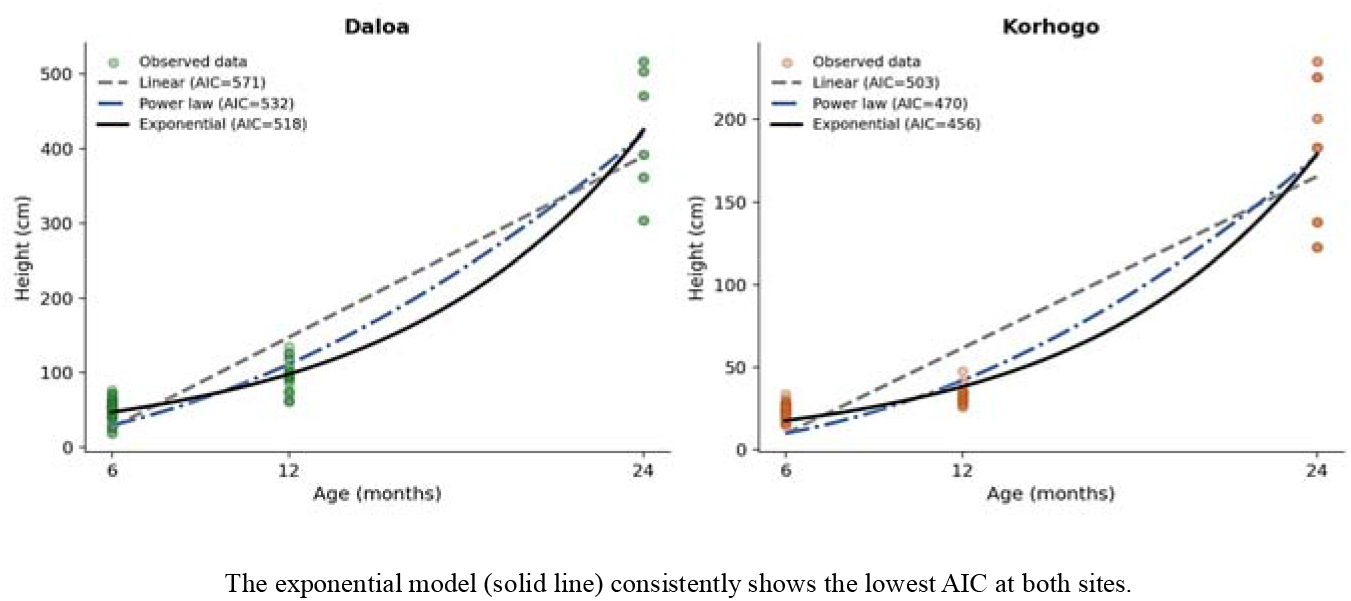
Comparison of the three growth models (linear, power law, exponential) fitted to individual height data of *Khaya senegalensis*, by site.

#### Collar-diameter growth

Unlike height, the exponential model is not consistently the best fit for collar diameter: at Daloa, the power law holds a slight advantage (Δ*AIC*=2.6 relative to the exponential; *D*=0.164×*age*^1.94); at Korhogo, the linear model narrowly wins (Δ*AIC*=0.7 relative to the power law; *D*=0.54+0.90×*age*), with the exponential fitting distinctly worse (Δ*AIC*=14.7). This divergence between height and diameter suggests that height growth in *Khaya senegalensis* remains, over this 6-24-month window, in a phase of continuous acceleration, whereas radial growth may already be entering a levelling-off phase, particularly at Korhogo (Table 8, Figure 10).

**Fig. 10.**
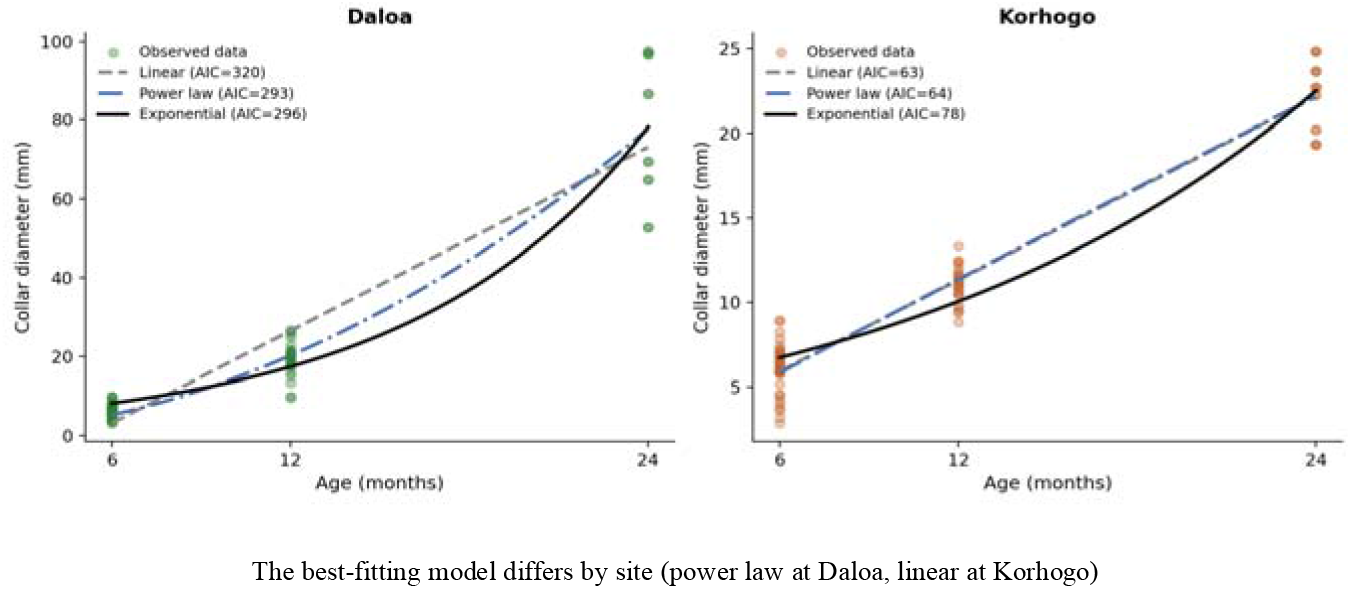
Comparison of the three growth models (linear, power law, exponential) fitted to individual collar-diameter data of *Khaya senegalensis*, by site.

##### f) Between-site phenotypic plasticity

A between-site phenotypic plasticity index (*RDPI*, Relative Distance Plasticity Index) was calculated for height, by seed-tree group and age, based on the relative difference between the Daloa and Korhogo means (Figure 11). This index ranges from 0.23 to 0.62 depending on the group and age considered, but this variation is correlated neither with seed-tree DBH (*r*=-0.14; *p*=0.57, all data combined), nor significantly different among seed-tree groups (ANOVA: *F*=0.59; *p*=0.71). This result indicates that, while mean seedling performance varies markedly between sites, the extent of this variation (plasticity itself) likewise does not depend on the size of the original seed tree. A result that closes one last possible door to a hidden seed-tree effect.

**Fig. 11.**
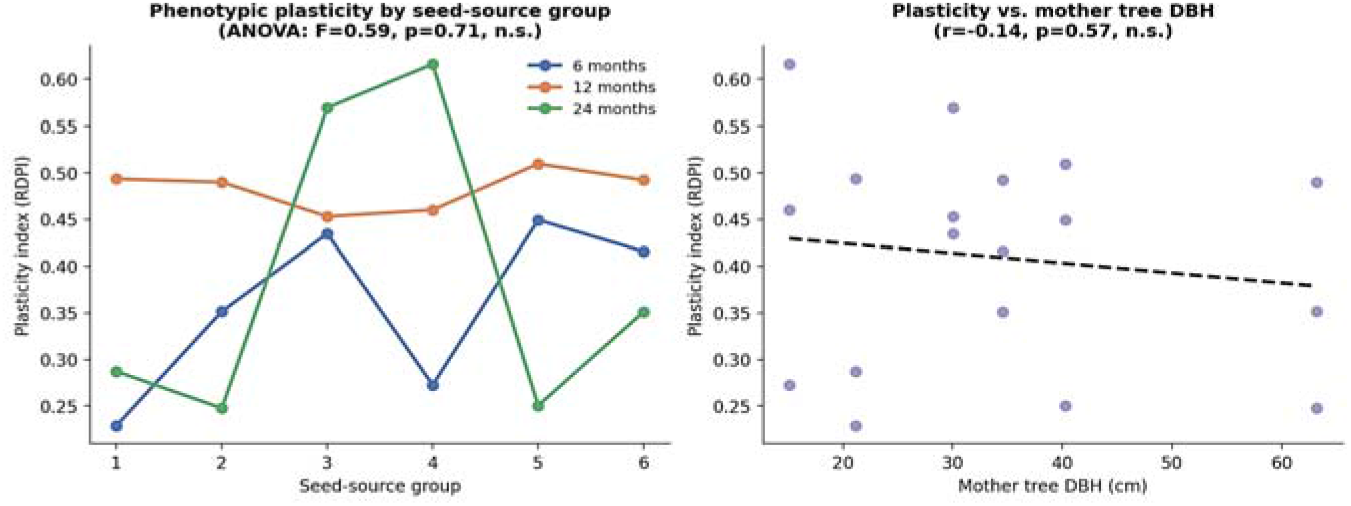
Between-site phenotypic plasticity index (*RDPI*) for height, by seed-source group and age, and its relationship with mother tree DBH

##### g) Allometric equations and model comparison

Allometric relationships between collar diameter and the various components of dry biomass are strong and highly significant (*R²* = 0.74 to 0.95; Figure 12), with clearly marked coefficients of determination depending on the organ considered. Total dry biomass follows a near-cubic power law of diameter (*exponent 2.58*), consistent with scaling laws typically observed in juvenile woody plants. Root biomass shows the strongest relationship with diameter (*R²*=0.92), making it a good non-destructive predictor of root investment, particularly useful for monitoring seedling establishment without systematic uprooting. The root-to-shoot ratio (root biomass relative to aboveground biomass) increases with age, rising from an average of 0.46 at 6 months to 1.39 at 12 months, reflecting a proportionally increasing root investment, a strategy consistent with the progressive acquisition of deep water resources in this deep-rooting species.

**Fig. 12.**
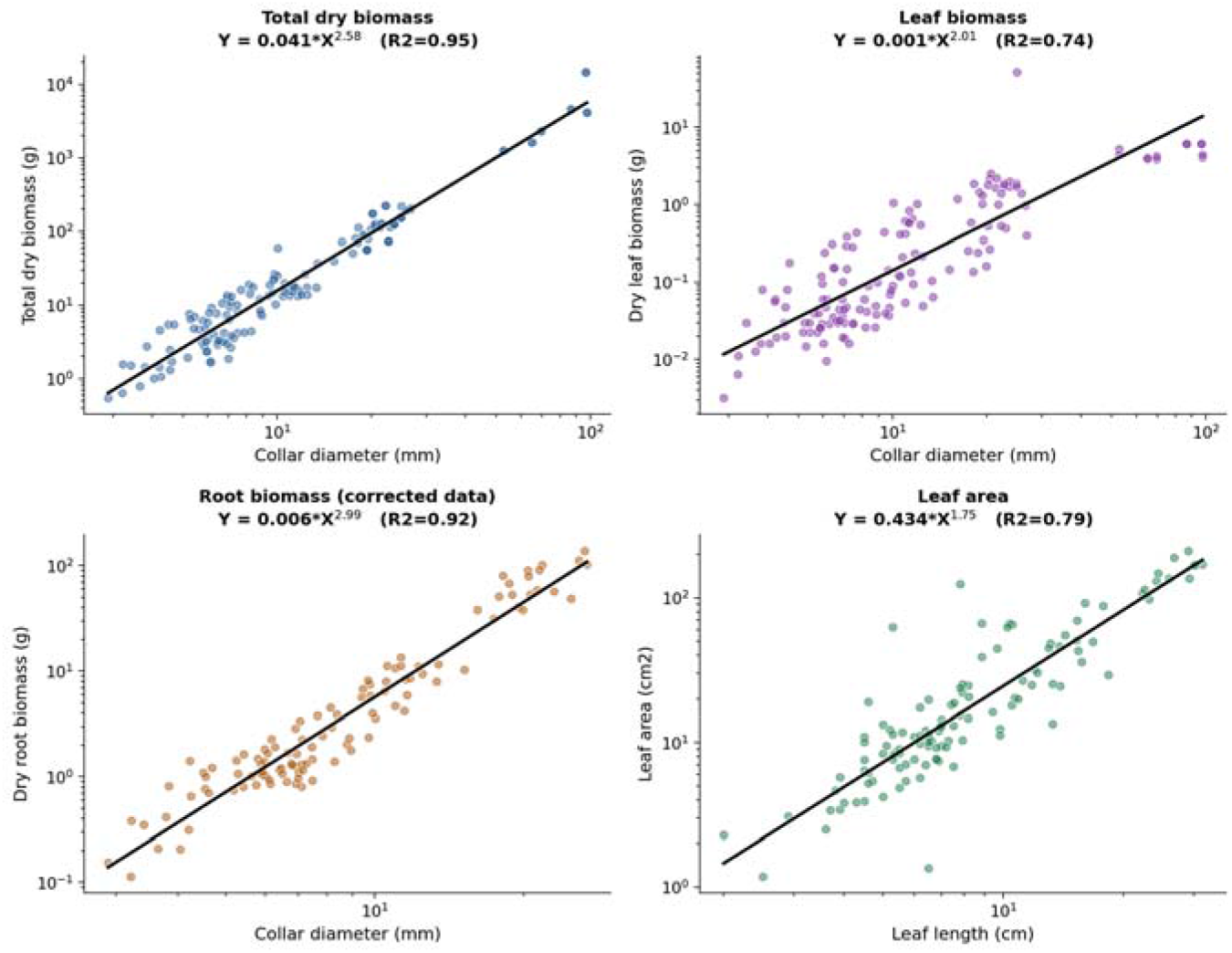
Allometric equations (log-log regression) relating collar diameter or leaf length to various biomass components and leaf area in *Khaya senegalensis* seedlings (6-24 months, sites pooled).

A single-predictor allometric model (total biomass ∼ diameter alone) was then compared with a two-predictor model (diameter and height), by *AIC* calculated on a logarithmic scale (Table 9). The two-predictor model is markedly superior (Δ*AIC*=97 relative to the single-predictor model), with a substantial reduction in residual error (*R²* raised to 0.974). The retained equation is: *ln(Biomass)* = -4.50 + 1.73×*ln(Diameter)* + 0.83×*ln(Height)*, providing a substantially more accurate non-destructive estimation tool than the single-predictor equation for nursery and reforestation-trial monitoring.

**Table 9.** Comparison of allometric models of total dry biomass, by *AIC* (logarithmic scale)

| Model | AIC | $R^2$ |
| --- | --- | --- |
| Power law (diameter only) | 206.1 | 0.948 |
| Quadratic log-log (diameter only) | 204.8 | 0.949 |
| Multiple allometry (diameter + height) | 109.2 | 0.974 |

##### h) Correlation structure of morphological traits

The correlation matrix (Figure 13) reveals strong phenotypic integration among growth traits (height, collar diameter, leaf area, and total biomass), consistent with coordinated juvenile growth across organs. Height, collar diameter, and leaf area are strongly correlated with one another (*r*>0.90), as well as with total and root biomass. Root length shows more moderate correlations with aboveground traits (*r*=0.65-0.87), suggesting a degree of independence in allocation to fine roots relative to height growth.

**Fig. 13.**
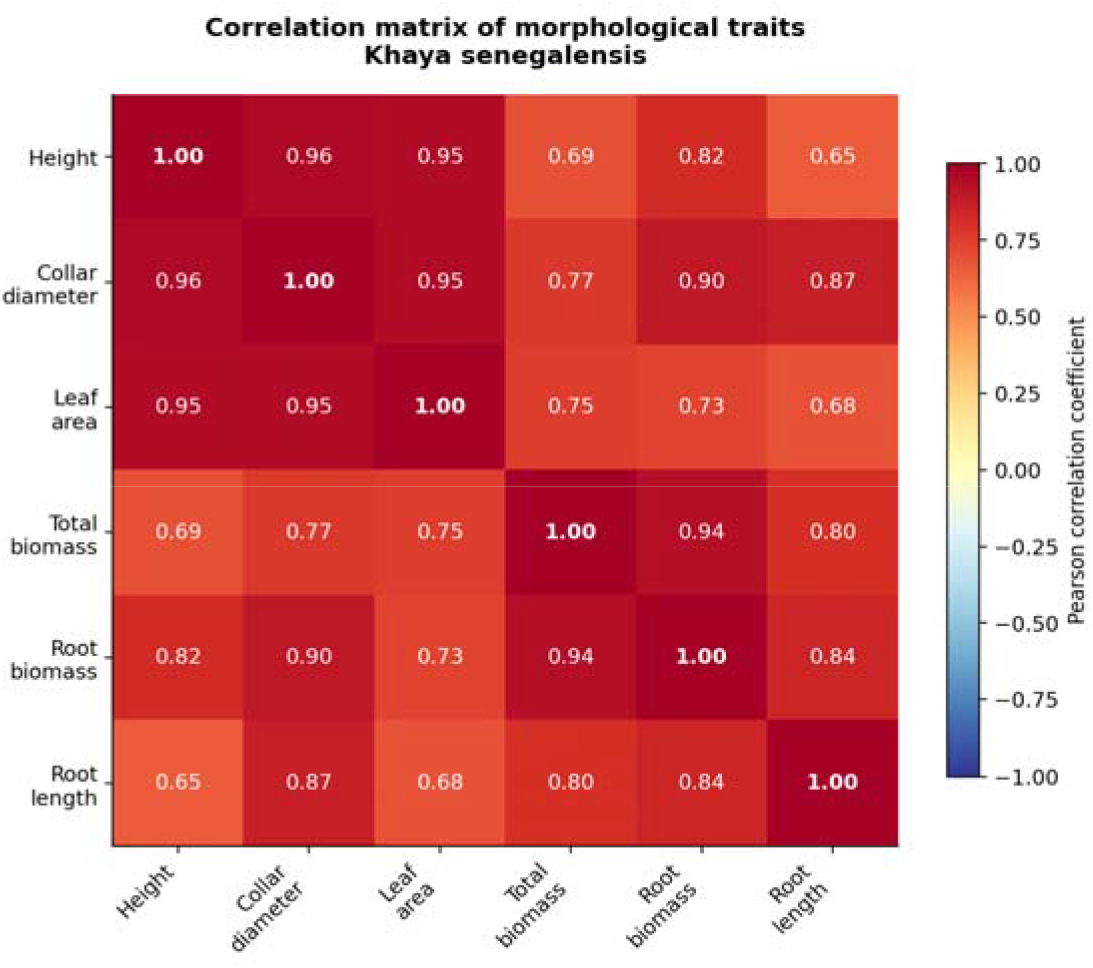
Pearson correlation matrix of the main morphological traits measured on *Khaya senegalensis* seedlings.

## 4. Discussion

### 4.1. Site, not seed tree, determines the regeneration of Khaya senegalensis, from 4 to 24 months

The results obtained here, converging across six independent analytical approaches (classical analysis of variance, box plots, continuous regression, mixed model with interaction, and this from the 4-month stage through to 24 months), unambiguously confirm hypothesis H1 and refute the hypothesis of a direct seed-tree effect (H2): the planting site exerts a decisive influence on germination and on the entire juvenile growth trajectory of *K. senegalensis*, whereas no dendrometric characteristic of the seed tree affects the performance of its offspring, at any measurement age. This result extends and reinforces that of Adji et al. (2020), who had already observed, in a design comparing three contrasting environments, a significant site effect but no effect of the six sampled seed trees on nearly all germination and development parameters. However, the present study goes further both methodologically and temporally: whereas those authors compared discrete seed-tree categories by analysis of variance at a single age, we directly regressed performance parameters on the actual diameter and height of the mother trees, confirmed the absence of an effect with a mixed model explicitly controlling for the non-independence of individuals from the same seed tree (Bates et al., 2015), and extended this test to four age classes (4, 6, 12, and 24 months). The convergence of these approaches across the entire juvenile developmental window, from the emergence of the first compound leaves through to the second year of growth, constitutes evidence considerably more robust and temporally extensive than an analysis confined to a single age.

A notable finding from this extension to the 4-month stage is the persistence of a contrast between traits: as at 6-24 months, height, collar diameter, number of leaves, and internode length do not differ according to seed-tree group, whereas leaf length and width differ significantly. This exception localised to leaf dimensions, already noted by the same authors at a later stage, could reflect an early plastic component of leaf area, more sensitive to small variations in germination conditions than structural traits (height, diameter), without thereby calling into question the overall conclusion of an absence of seed-tree effect on the offspring’s overall performance.

The classical ANOVA with main effects (site, seed-tree group, age), conducted in addition to the continuous-regression and mixed-model approaches to reproduce the analytical format usually employed in this type of study (Adji et al., 2020; Ky-Dembele et al., 2014), fully confirms these results: site and age are highly significant for all three growth responses tested, whereas seed-tree group never is. The fact that a more classical analytical method, more familiar to readers of this type of literature, arrives at the same conclusion as the more elaborate approaches reinforces the credibility of the result among an agronomic readership accustomed to this format, while demonstrating that the conclusion does not depend on any particular methodological choice.

### 4.2. A seed carrying no seed-tree size effect: where, then, does the maternal effect go?

The mediation analysis, explicitly testing hypothesis H3 of an indirect seed-tree effect via seed mass, closes one last possible door to a hidden maternal effect: the first link in the causal chain (the ability of seed-tree DBH to predict the mass of the seeds it produces) is not significant (*r*=0.36; *p*=0.48), which structurally breaks any possibility of mediation, regardless of the strength of the subsequent links. This result should be viewed in light of the classical literature on maternal effects in plants, which highlights that these effects, although frequent and often powerful, are not systematically correlated with the size of the mother plant itself, but may instead depend on more specific factors such as reproductive age, the position of the fruit within the crown, or local microclimatic conditions during flowering and fruiting (Roach and Wulff, 1987). The phenological description of *K. senegalensis* germination (hypogeal, with a long epicotyl and nearly sessile prophylls prior to the establishment of alternate-spiral phyllotaxis) further illustrates early development strongly dependent on seed reserves rather than on inherited maternal-tree characteristics, consistent with the observed absence of a link between seed-tree size and seed mass.

It is also worth noting that Norden et al. (2009), in their comparative analysis of more than 1.000 tropical tree species, showed that seed mass is an excellent predictor of germination time at the interspecific level. This result also holds at the intraspecific level in *K. senegalensis* in our own mediation analysis (a positive trend between seed mass and germination rate, *r*=0.75, although not significant at the conventional threshold with only six seed-tree groups). The break therefore occurs precisely between tree size and the mass of the seed it produces, and not between seed mass and germination performance. A distinction that, to our knowledge, had never before been tested so explicitly in this species.

### 4.3. Exponential growth in height, but earlier levelling-off of diameter

The comparison of non-linear growth models indicates, for height, an exponential rather than linear or power-law regime at both sites (Δ*AIC* = 13.6 at Korhogo and 14.7 at Daloa relative to the power-law model; Burnham and Anderson, 2002b, 2002a). This result warrants cautious interpretation: an exponential fit over a time window of only 24 months cannot distinguish genuine sustained exponential growth from an early phase of a broader sigmoidal model, whose inflection point and asymptote would only become perceptible at a more advanced age. This constitutes a limitation that we explicitly acknowledged by excluding the fit of a Gompertz model, whose asymptote parameters proved statistically poorly constrained by the available data. This finding aligns with methodological recommendations in the literature on fitting non-linear growth models in plant ecology, which stress the need for observations covering the entire life trajectory to reliably estimate the parameters of an asymptotic model (Paine et al., 2012).

Collar diameter, by contrast, follows a more nuanced pattern; the power law prevailing at Daloa and the linear model at Korhogo; the exponential being in both cases the worst-fitting model. This divergence between height and diameter suggests that radial growth in *K. senegalensis* enters a levelling-off phase earlier than height growth. This is a pattern of differential allocation consistent with the strategy of many juvenile woody species that prioritise vertical growth (access to light) before investing further in radial thickening. Extended monitoring beyond 24 months, particularly of the 5-and 7-year-old individuals already available at Daloa within the broader design of this study, would allow this hypothesis of a differential timeline between the two growth axes to be directly tested.

### 4.4. Phenotypic plasticity and allometric equations: two complementary tools for management

The between-site phenotypic plasticity index, although variable from one seed-tree group to another, is correlated neither with seed-tree DBH nor with group membership. This is indeed a result that harmoniously complements the study’s central finding: not only does seed-tree size fail to affect the mean performance of its offspring, but it also fails to affect the offspring’s sensitivity to site conditions. This lack of structuring of plasticity by maternal origin suggests that the response of *K. senegalensis* to the bioclimatic gradient tested here is a property shared across the entire population studied, consistent with the hypothesis of broad ecological tolerance in this species already noted in the literature on its genetic diversity (Bouka Dipelet et al., 2019).

On a more applied level, the comparison of allometric models demonstrates the value of systematically incorporating height, in addition to collar diameter, into biomass prediction equations (Δ*AIC* = 97 in favour of the two-predictor model; *R²* raised from 0.948 to 0.974). This result is consistent with the reference literature on tropical tree allometry, which has established, since the work of Chave et al. (2014) that incorporating total height substantially improves the accuracy of biomass models compared with diameter alone. The increase in the root-to-shoot ratio between 6 and 12 months (from 0.46 to 1.39) further reflects a proportionally increasing root investment, consistent with the progressive acquisition of deep water resources in this deep-rooting species, characteristic of Sudano-Guinean savanna tree species.

### 4.5. Implications for conservation and study limitations

Taken together, these results call for a pragmatic reorientation of priorities in restoration and agroforestry programmes based on *K. senegalensis*: rather than investing resources in fine-grained selection of seed trees based on size criteria, managers would benefit from focusing their efforts on the judicious choice of planting site, an approach increasingly formalised in the climate-smart restoration tools developed for tropical tree species (Fremout et al., 2022; St.Clair et al., 2022).

This study has several limitations. Data quality control revealed physically inconsistent root-biomass values (5.9% of individuals), treated as missing rather than arbitrarily corrected. The number of seed-tree groups (six) inherently limits the power of the mediation analysis. The monitoring window (24 months for the analyses presented) does not allow a definitive conclusion on the complete shape of the species’ growth trajectory, which can live for several decades. Monitoring at 5 and 7 years, already under way at Daloa, will eventually resolve this ambiguity. Finally, this study was conducted at two sites: replication across a larger number of sites would allow the conclusion to be generalised to the species’ entire Sudano-Guinean distribution range.

## 5. Conclusion

Faced with the urgent need to restore *Khaya senegalensis* stands, a Vulnerable species whose natural regeneration is collapsing under the combined pressure of overexploitation and climate change. This study provides a clear answer to a question that has hitherto lacked solid statistical proof: it is not seed-tree size but the choice of planting site that determines the success of germination and juvenile growth in this species, from the earliest months of development through to two years of age. Converging across six independent approaches (from classical analysis of variance to formal mediation), this result methodically closes off every avenue through which a seed-tree effect might have manifested. The multi-predictor allometric equations developed here also offer a non-destructive monitoring tool immediately applicable in nurseries. Ultimately, this study invites a refocusing of savanna mahogany restoration programmes on a simpler, yet more decisive, question than seed-tree selection: *where to plant, rather than which seeds to choose*.

## CRediT authorship contribution statement

**Beda Innocent Adji**: Conceptualization, Data curation, Formal analysis, Investigation, Methodology, Resources, Software, Validation, Visualization, Writing – original draft, Writing - review & editing. **Doffou Selastique Akaffou**: Supervision, Validation.

## Acknowledgements

The authors are deeply grateful to the rural populations of the various localities (Katiola, Niakara, Korhogo, and Sinématiali) visited for their mobilisation, generosity, and hospitality during the phases of seed provenance collection and mother-tree sampling. The authors thank the various owners of the agroforestry parklands visited for making their orchards available and granting permission to conduct this study on the various trees within them. Finally, the authors thank the *Centre National de Recherche Agronomique* of Côte d’Ivoire (CNRA) for making available the Diabaté Kamonon forestry research station in Sédiakaha, Korhogo, for part of this study.

## Data availability

The datasets analysed as part of this study are available at: ADJI, B. I. (2026). Supporting Dataset for the document of Germination, morphology and biomass of *Khaya senegalensis*, *Pterocarpus erinaceus* and *Parkia biglobosa* seedlings according to mother-tree dendrometric characteristics and planting site in Côte d’Ivoire (2018-2025). https://doi.org/10.5281/zenodo.21435509

